# Bioinformatics Copilot 1.0: A Large Language Model-powered Software for the Analysis of Transcriptomic Data

**DOI:** 10.1101/2024.04.11.588958

**Authors:** Yongheng Wang, Weidi Zhang, Siyu Lin, Matthew S. Farruggio, Aijun Wang

## Abstract

The field of single-cell transcriptomics has been producing extensive datasets, advancing our understanding of cellular functions in various tissues, and empowering diagnosis, prognosis, and drug development. However, parsing through this data has been a monumental task, often stretching weeks to months. This bottleneck arises due to the sheer volume of data generated—ranging from hundreds of gigabytes to tens of terabytes—that demands extensive time for analysis. Moreover, the data analysis involves an intricate series of steps utilizing various software packages, creating a steep learning curve for biologists. Additionally, the iterative nature of data analysis in this domain necessitates a deep biological insight to formulate relevant questions, conduct analysis, interpret results, and refine hypotheses. This iterative loop has required close collaboration between biologists and bioinformaticians, which is hampered by protracted communication cycles. To address these challenges, we present a large language model-powered software, Bioinformatics Copilot 1.0. It allows users to analyze data through an intuitive natural language interface, without requiring proficiency in programming languages such as Python or R. It is engineered for cross-platform functionality, with support for Mac, Windows, and Linux. Importantly, it facilitates local data analysis, ensuring adherence to stringent data management regulations that govern the use of patient samples in medical and research institutions. We anticipate that this tool will expedite the data analysis workflow in numerous research endeavors, thereby accelerating advancements in the biomedical sciences.

## Introduction

Since 2009, a variety of single-cell sequencing technologies relying on next-generation sequencing have advanced rapidly. Among these, spatially resolved transcriptomics, a type of single-cell omics technology, has shown remarkable promise and has been recognized as the “Method of the Year 2020” by Nature Methods^1^. Such technologies allow for the visualization and quantification of thousands of transcripts with sub-cellular resolution, demonstrating significant advantages over traditional methods (e.g., western blot, immunohistochemistry staining), where one or a few genes are assessed in one experiment. Thus, these technologies have been used to study cancer^2,3^, embryo development^4^, and neuroscience^5,6^ to revisit important questions and reveal unprecedented insights. As more technologies become commercially available, it is anticipated that they will be harnessed to set new standards for biological research, diagnostics, and prognosis. Nevertheless, biomedical scientists face challenges, notably the size of datasets, which are often vast and can take months to years to analyze. Despite the development of over 1,000 tools in the past decade to aid in single-cell data analysis^7,8^, data processing remains a bottleneck for many projects. Additionally, a significant issue is that many biologists lack training in coding languages such as R or Python, which hampers their ability to effectively analyze data.

In the past two decades, artificial Intelligence has seen a remarkable rise in prominence, propelled by monumental strides in computing power, data storage capacities, and the gathering of extensive datasets. This era has been defined by the advent of intricate machine learning algorithms, highlighted by the debut of transformer models^9^ and the advent of platforms such as ChatGPT. Such breakthroughs have led to the development of applications utilizing Large Language Models (LLMs)^10^ in diverse areas like text, voice, image, and video creation. These innovations have significantly improved the efficiency of design, production, writing, and filmmaking. With the widespread adoption of large language models, researchers have begun employing natural language prompts to simplify coding tasks, particularly in data analysis and code completion^11^. Consequently, these innovations have paved the way for the development of a specialized tool, Bioinformatics Copilot, aimed at expediting bioinformatics analysis through the integration of LLMs.

## Result

The Bioinformatics Copilot user interface is sectioned into five distinct areas (Fig. 1). At the top, there is a designated space where users can upload their files for analysis. On the bottom left, there is a space where users can input their prompts. On the top left, the copilot will chat with the users in natural language. On the top right, the copilot will generate code and report the progress. On the bottom right, the copilot will display the figures.

**Fig. 1.**
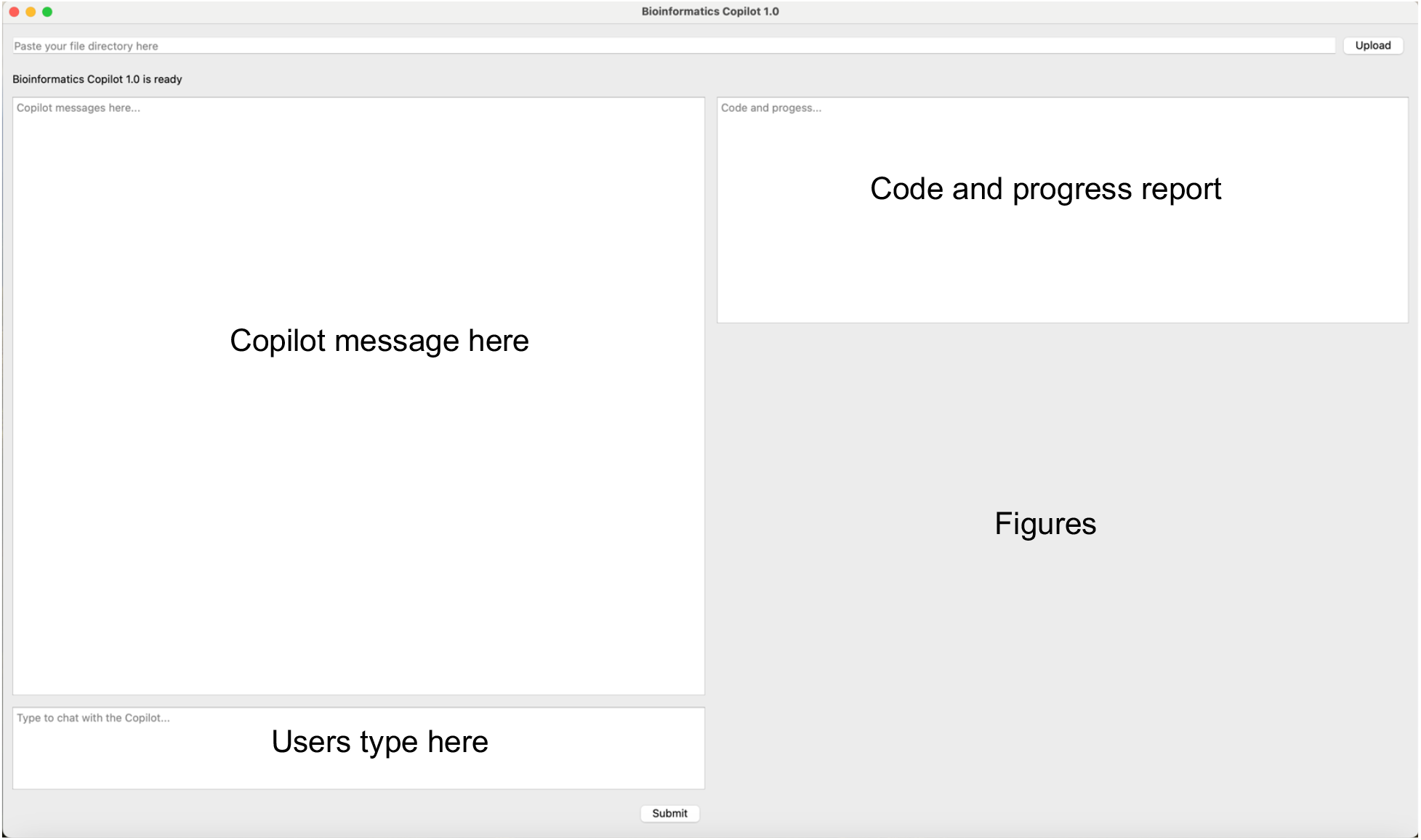
The user interface of Bioinformatics Copilot.

Human cells have more than 20,000 genes. The expression level of these genes is frequently linked to the likelihood of developing cancer and can serve as a measure of how effective a treatment might be. Therefore, assessing gene expression levels with high-throughput methods can assist scientists in thoroughly evaluating the effectiveness of drugs or treatments. To assess the expression level and location of a specific gene, such as MALAT1, the user might input, “Can you make a feature plot for MALAT1?” The copilot responds by creating the visualization using the ImageFeaturePlot function (Fig. 2).

**Fig. 2.**
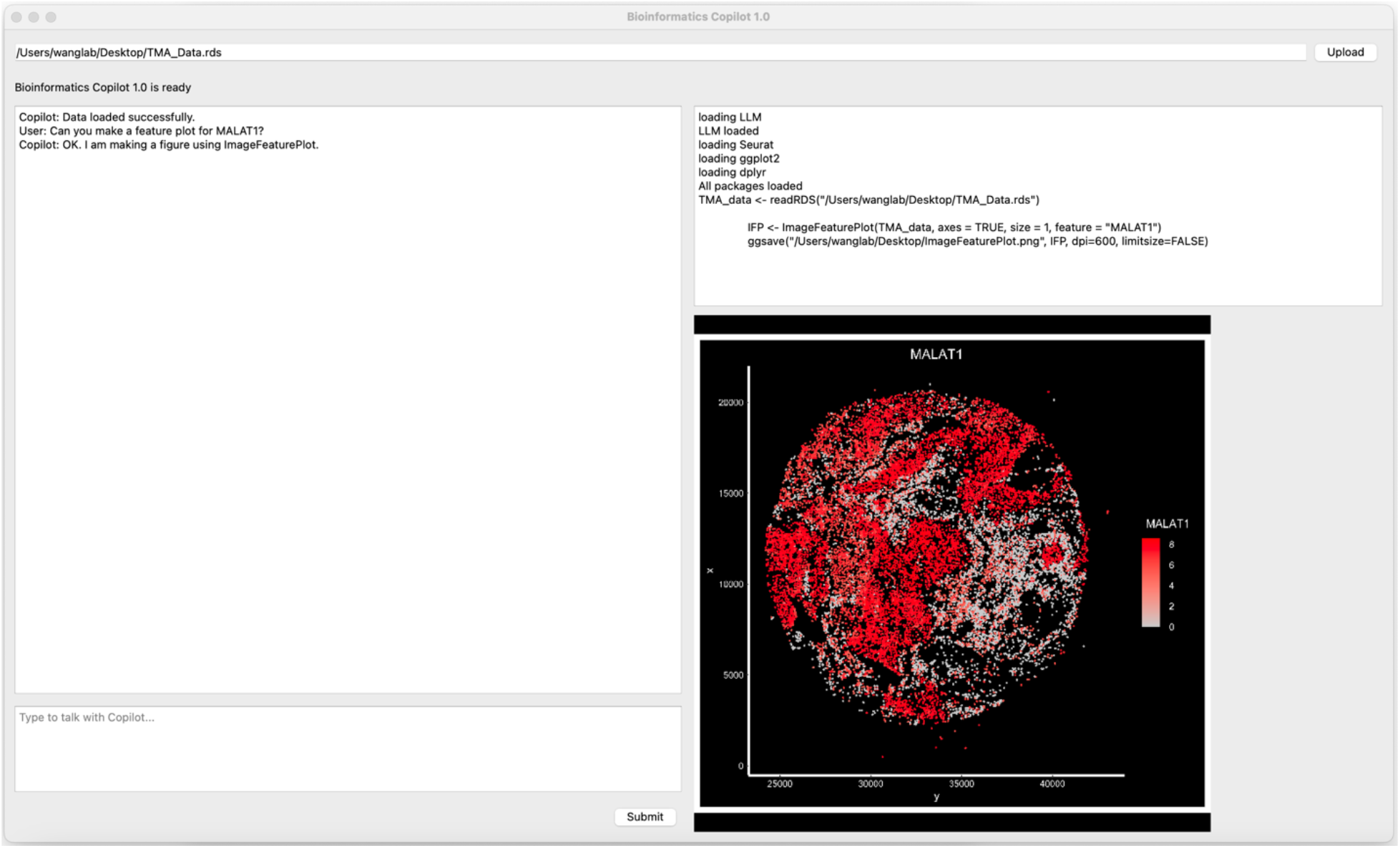
Evaluation of gene expression using Bioinformatics Copilot.

Every type of tissue consists of a diversity of cell types, each fulfilling unique roles. Take tumor tissue as an example: it encompasses a range of cells such as macrophages, T cells, tumor cells, and the cells that form blood vessels. Charting the cellular composition of these tissues is vital for gauging their state. Through such analysis, a doctor can determine whether a treatment like an anti-angiogenesis drug has successfully obstructed the development of new blood vessels or verify whether chimeric antigen receptor (CAR) T cells have effectively infiltrated the tumor. To evaluate the cell composition of a tissue, users can inquire about the Uniform Manifold Approximation and Projection (UMAP) for a given tissue sample. By simply formulating the request in natural language, the copilot will generate a figure revealing the different clusters of cells in the tissue (Fig.3).

**Fig. 3.**
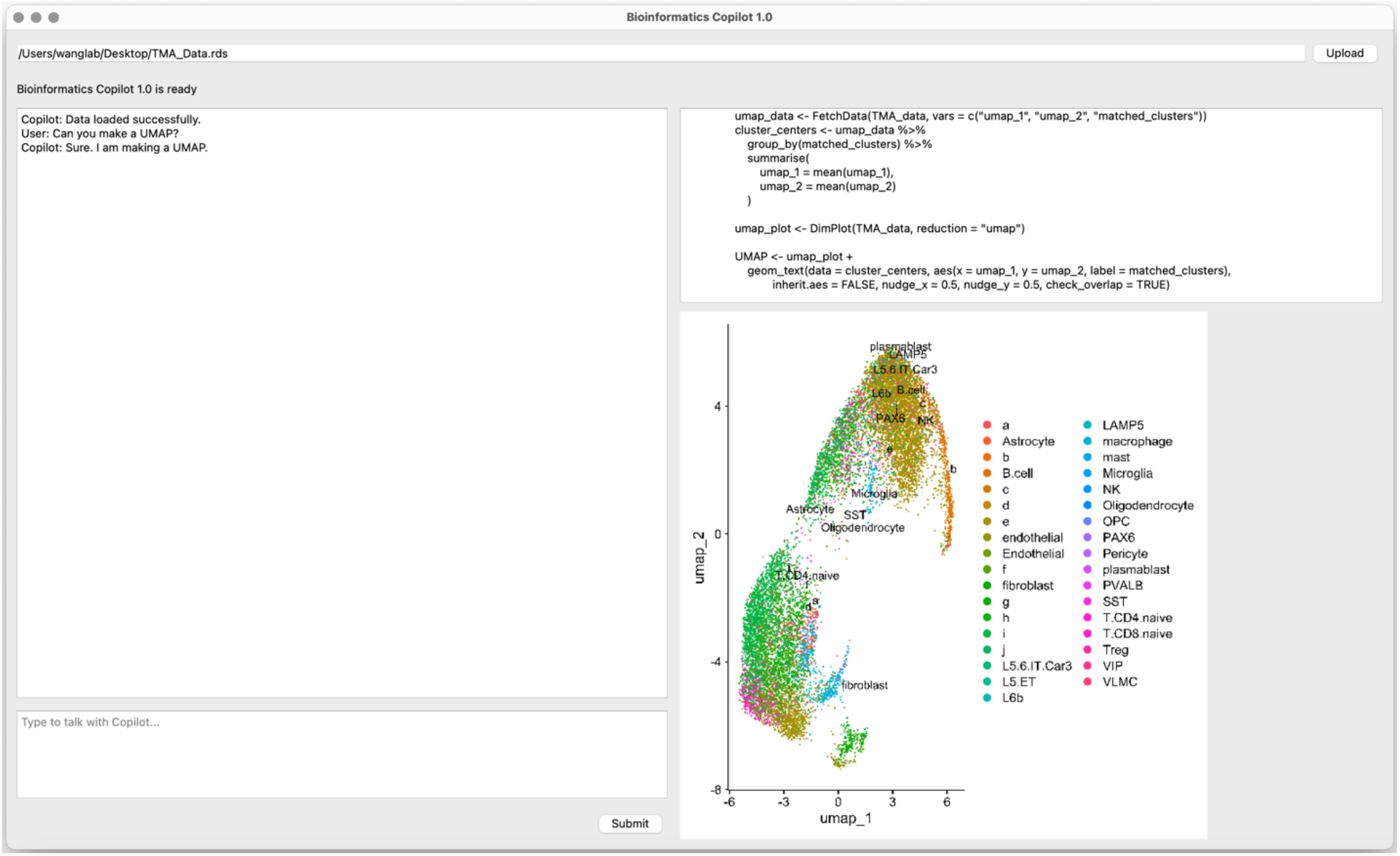
Assessment of cell clustering using Bioinformatics Copilot.

Mapping the precise location of cells in a tissue is crucial for understanding a lot of biological phenomena. For example, in the context of tumor tissue, one could examine the variance in gene expression between CAR T cells that are in direct contact with tumor cells and those that are not. Such an analysis could unveil insights into the molecular mechanisms that govern cancer cell recognition, T cell exhaustion, etc. In brain tissue, diverse cell types such as neurons, oligodendrocytes, and astrocytes engage in functions including metabolic and structural support, as well as signal transmission. Knowledge of cell spatial distribution in neurological diseases can be pivotal in dissecting the disease’s origin and progression, aiding in the development of targeted interventions. To visualize the spatial information of the different cell types, the user can type “Can you make a dimension plot?” The copilot will generate a figure showing the location of the different cells in the tissue (Fig.4).

**Fig. 4.**
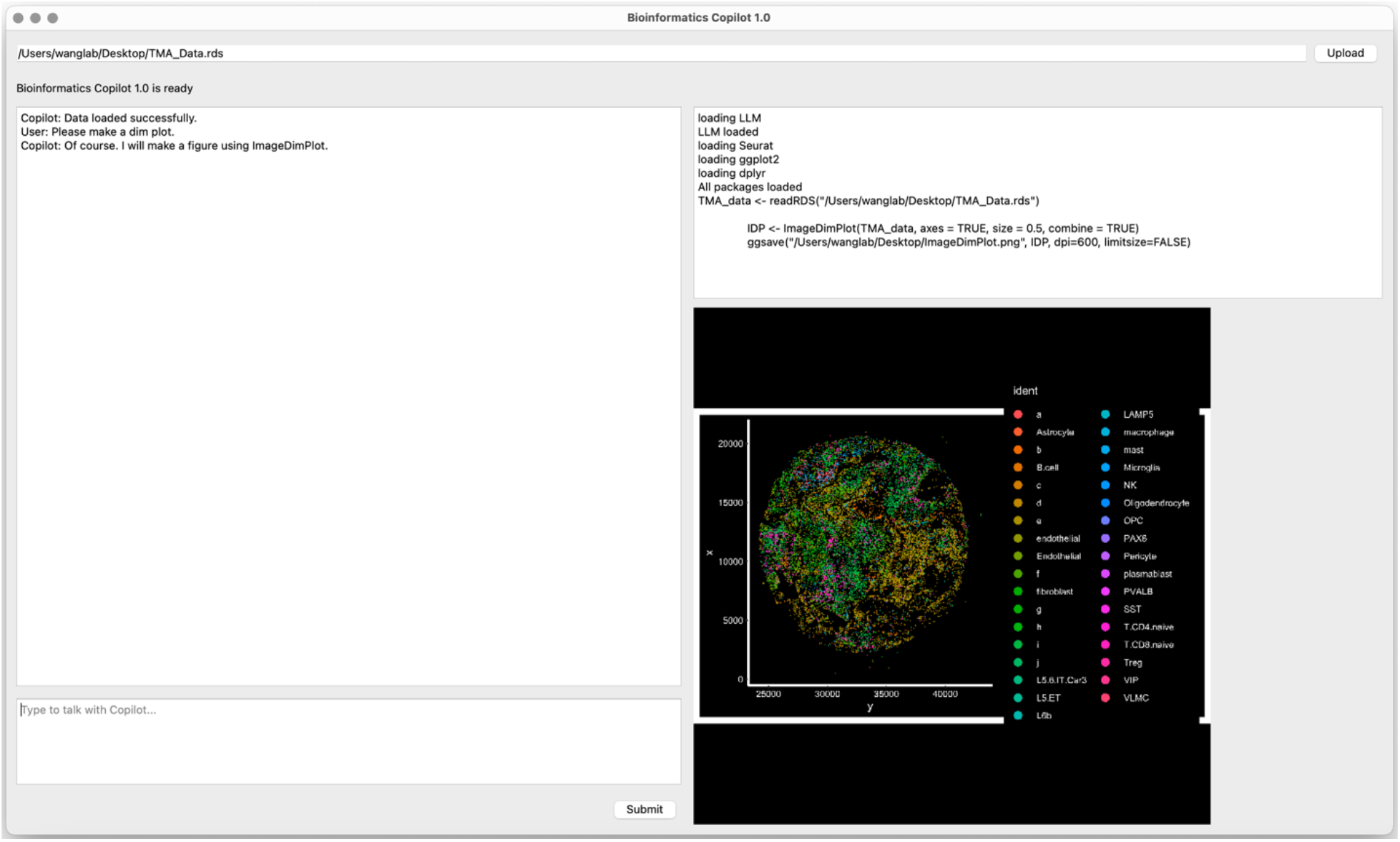
Visualization of the locations of different cell types.

## Discussion

The current version of the software is only equipped to analyze Seurat objects. At present, it is not capable of directly handling larger FASTQ datasets, which can often reach the size of several hundred gigabytes or even extend to terabytes. Consequently, users must procure Seurat objects from either bioinformatics core facilities or external service providers. The objective of this version is to streamline the oftentimes lengthy communication process between bench scientists and bioinformatics experts. To further assist biologists in data analysis, future updates will include features for performing genome mapping on a local server via intuitive natural language. This advancement will allow biologists to proceed with data analysis immediately following the sequencing step, thereby expediting the progress of their projects. In addition, the introduction of multi-agent and “self-operating computer” has the potential to unlock new possibilities and improve software performance.

## Conclusion

We have developed LLM-powered software, Bioinformatics Copilot 1.0, for analyzing transcriptomic data using natural language. This tool is anticipated to accelerate a multitude of research endeavors and help advance the field of biomedical sciences.

## Materials and Methods

Detailed methods will be released in the peer-reviewed article. Briefly, LLM is deployed to understand the prompt and output information needed for executing specific tasks with R, such as generating plots (ImageDimPlot, ImageFeaturePlot, UMAP, etc.).

## References

1. Method of the Year 2020: spatially resolved transcriptomics. (2021). Nat Methods 18, 1. 10.1038/s41592-020-01042-x.

2. Janesick, A., Shelansky, R., Gottscho, A.D., Wagner, F., Williams, S.R., Rouault, M., Beliakoff, G., Morrison, C.A., Oliveira, M.F., Sicherman, J.T., et al. (2023). High resolution mapping of the tumor microenvironment using integrated single-cell, spatial and in situ analysis. Nat Commun 14, 8353. 10.1038/s41467-023-43458-x.

3. He, S., Bhatt, R., Brown, C., Brown, E.A., Buhr, D.L., Chantranuvatana, K., Danaher, P., Dunaway, D., Garrison, R.G., Geiss, G., et al. (2022). High-plex imaging of RNA and proteins at subcellular resolution in fixed tissue by spatial molecular imaging. Nat Biotechnol 40, 1794–1806. 10.1038/s41587-022-01483-z.

4. Chen, A., Liao, S., Cheng, M., Ma, K., Wu, L., Lai, Y., Qiu, X., Yang, J., Xu, J., Hao, S., et al. (2022). Spatiotemporal transcriptomic atlas of mouse organogenesis using DNA nanoball-patterned arrays. Cell 185, 1777–1792 e1721. 10.1016/j.cell.2022.04.003.

5. Yao, Z., van Velthoven, C.T.J., Kunst, M., Zhang, M., McMillen, D., Lee, C., Jung, W., Goldy, J., Abdelhak, A., Aitken, M., et al. (2023). A high-resolution transcriptomic and spatial atlas of cell types in the whole mouse brain. Nature 624, 317–332. 10.1038/s41586-023-06812-z.

6. Zhang, M., Pan, X., Jung, W., Halpern, A.R., Eichhorn, S.W., Lei, Z., Cohen, L., Smith, K.A., Tasic, B., Yao, Z., et al. (2023). Molecularly defined and spatially resolved cell atlas of the whole mouse brain. Nature 624, 343–354. 10.1038/s41586-023-06808-9.

7. Zappia, L., and Theis, F.J. (2021). Over 1000 tools reveal trends in the single-cell RNA-seq analysis landscape. Genome Biol 22, 301. 10.1186/s13059-021-02519-4.

8. Dance, A. (2022). Which single-cell analysis tool is best? Scientists offer advice. Nature 612, 577–579. 10.1038/d41586-022-04426-5.

9. Vaswani, A., Shazeer, N., Parmar, N., Uszkoreit, J., Jones, L., Gomez, A. N., Kaiser, L., & Polosukhin, I. (2017). Attention is all you need. arXiv preprint arXiv:1706.03762v7. 10.48550/arXiv.1706.03762.

10. Naveed, H., Khan, A. U., Qiu, S., Saqib, M., Anwar, S., Usman, M., Akhtar, N., Barnes, N., & Mian, A. (2023). A Comprehensive Overview of Large Language Models. arXiv preprint arXiv:2307.06435, last revised 9 April 2024. 10.48550/arXiv.2307.06435.

11. Mastropaolo, A., Pascarella, L., Guglielmi, E., Ciniselli, M., Scalabrino, S., Oliveto, R., Bavota, G. (2023). On the Robustness of Code Generation Techniques: An Empirical Study on GitHub Copilot. arXiv preprint arXiv:2302.00438. 10.48550/arXiv.2302.00438

